# scAI-SNP: a method for inferring ancestry from single-cell data

**DOI:** 10.1101/2024.05.14.594208

**Authors:** Sung Chul Hong, Francesc Muyas, Isidro Cortés-Ciriano, Sahand Hormoz

## Abstract

Collaborative efforts, such as the Human Cell Atlas, are rapidly accumulating large amounts of single-cell data. To ensure that single-cell atlases are representative of human genetic diversity, we need to determine the ancestry of the donors from whom single-cell data are generated. Self-reporting of race and ethnicity, although important, can be biased and is not always available for the datasets already collected. Here, we introduce scAI-SNP, a tool to infer ancestry directly from single-cell genomics data. To train scAI-SNP, we identified 4.5 million ancestry-informative single-nucleotide polymorphisms (SNPs) in the 1000 Genomes Project dataset across 3201 individuals from 26 population groups. For a query single-cell data set, scAI-SNP uses these ancestry-informative SNPs to compute the contribution of each of the 26 population groups to the ancestry of the donor from whom the cells were obtained. Using diverse single-cell data sets with matched whole-genome sequencing data, we show that scAI-SNP is robust to the sparsity of single-cell data, can accurately and consistently infer ancestry from samples derived from diverse types of tissues and cancer cells, and can be applied to different modalities of single-cell profiling assays, such as single-cell RNA-seq and single-cell ATAC-seq. Finally, we argue that ensuring that single-cell atlases represent diverse ancestry, ideally alongside race and ethnicity, is ultimately important for improved and equitable health outcomes by accounting for human diversity.

## 1. Introduction

In recent years, single-cell technologies, especially single-cell RNA sequencing (scRNA-seq) and single-cell transposase-accessible chromatin sequencing (scATAC-seq), have become ubiquitous, revolutionizing our understanding of cellular heterogeneity in many biological systems. Consequently, many datasets have emerged, epitomized by ambitious endeavors such as the Human Cell Atlas^1,2^ and the Human Tumor Atlas Network^3^, which aim to create comprehensive reference maps of all human cells in healthy and diseased tissues. These data sets hold the potential for groundbreaking discoveries in personalized medicine, developmental biology, and a myriad of other fields.

The utility and impact of a reference atlas, such as the Human Cell Atlas, hinge crucially on its representativeness of human diversity. It is imperative to construct the reference atlases from a wide array of donors, encompassing all ancestries. This diversity is not merely a demographic criterion but a fundamental biological necessity. Genetic variation across different populations can significantly influence gene expression patterns, epigenetic landscapes, and cellular responses to environmental stimuli^4–6^. Without a diverse representation, reference atlases risk harboring biases, potentially leading to erroneous biological conclusions and limiting their applicability in global health contexts. For example, hematological characterization of individuals of European descent has shown significant differences in certain blood counts compared with those of African descent^7^. Consequently, reference datasets of blood counts constructed from predominantly European individuals have misled physicians into prescribing inappropriately low dosages of chemotherapy for early-stage breast cancer patients of African descent, who had different baseline ranges of neutrophils, resulting in poorer survival outcomes^8–10^. Therefore, single-cell atlases cannot truly represent the general population if the underlying data disproportionately represent or exclude specific ancestry groups.

One way to ensure that an atlas is representative of human genetic diversity is to require the donors to self-report their race or ethnicity. Unlike ancestry, which is determined by the genetic variants common to population groups that are inherited by an individual and can be reliably quantified through genetic analyses, race and ethnicity are socially constructed and ascribed. Race is socially assigned based on physical features, such as skin color, and ethnicity is socially assigned based on cultural similarities between individuals, such as a common language. However, using self-reported race or ethnicity to build a single-cell atlas has several drawbacks. People might not know or accurately report their complete racial or ethnic background, especially if they have a mixed heritage. Also, we cannot add self-reported race or ethnicity information retroactively to older samples that did not collect it. This limitation is particularly relevant given the vast repositories of historical single-cell data that hold potential insights. Furthermore, in genome-wide association studies (GWAS), using diverse groups based on ancestry in addition to race and ethnicity improves our ability to discover genetic predictors of complex traits^11^. Therefore, we need a method to infer ancestry directly from the single-cell data itself.

Single nucleotide polymorphisms (SNPs) that have significantly different frequencies across different population groups can be used to infer an individual’s ancestry. Some methods have focused on identifying a small subset of such SNPs that are most informative for determining population structures^12–14^. Although for some applications, such as microarray measurements, having a selective set of SNPs is cost-effective, it is more strategic with sparse data like scRNA-seq data to cast a wider net; that is, to leverage numerous sites in the genome. More recently, whole genome sequencing (WGS) has enabled the development of genome-wide approaches that account for all the SNPs detected in an individual’s genome when assigning ancestry. There are two broad approaches for inferring ancestry from whole genome sequences: global and local. In global inference, the SNPs are used to assign an individual to an ancestral group or a mixture of the ancestral groups^15–17^. In local inference, each region of the genome of an individual is assigned to an ancestral group or a mixture of such groups, and the goal is to find the appropriate boundaries of these regions^18^. Here, we focus on global inference of ancestry from single-cell data.

Two general approaches have been proposed for inferring global ancestry from sequencing data, parametric and non-parametric. Parametric approaches, such as STRUCTURE^16^, FRAPPE^19^, fastSTRUCTRE^20^, and ADMIXTURE^15^, find the optimal choice of parameters for a statistical model where the parameters correspond to the fraction that each ancestry group contributes to each individual in the data and the frequency of the alleles in each population group. The posterior distribution of the parameters can be found using Bayesian methods such as Markov-Chain Monte Carlo or approximated efficiently using variational methods that rely on optimization. Non-parametric approaches, such as EIGENSTRAT^17^, take a geometric perspective and identify the linear combinations of alleles that capture the directions of largest variation across the individuals in the data. The first few directions often capture the population structure in the data and can be used to assign the ancestral groups (or combinations thereof) to each individual^21^. Importantly, both approaches can work without any prior knowledge of the ancestral groups or their allele frequencies. The parametric methods can learn this information while simultaneously fitting a model to the data using expectation maximization, whereas non-parametric methods find the directions of maximum variation present in the data without relying on labeled individuals or population structure.

Our approach builds on the non-parametric methods for global inference of ancestry because these methods are more robust to missing the data and are fast. Specifically, similar to EIGENSTRAT, we use principal component analysis (PCA) to identify the most informative directions in allele space for inferring ancestry (see also^22^). Unlike EIGENSTRAT, our goal is not to only capture variation in a dataset due to population structure but also to use these directions to assign out-of-sample individuals to a known ancestry group. Therefore, we used a dataset of individuals with known (self-reported) ancestry from the 1000 Genomes Project (1KGP)^23^. We identified 4.5 million ancestry-informative SNPs in the 1KGP WGS dataset and converted each individual’s genotype to a 4.5 million-dimensional vector where each element of the vector corresponds to the allele of the given SNP site detected in that individual. Next, we conducted PCA on these vectors to find a low-dimensional representation for them (Figure 1b). Finally, we obtain a single vector representation of each population group by averaging the vector representation in PCA space of all individuals in the 1KGP data that belong to that population group.

**Figure 1.**
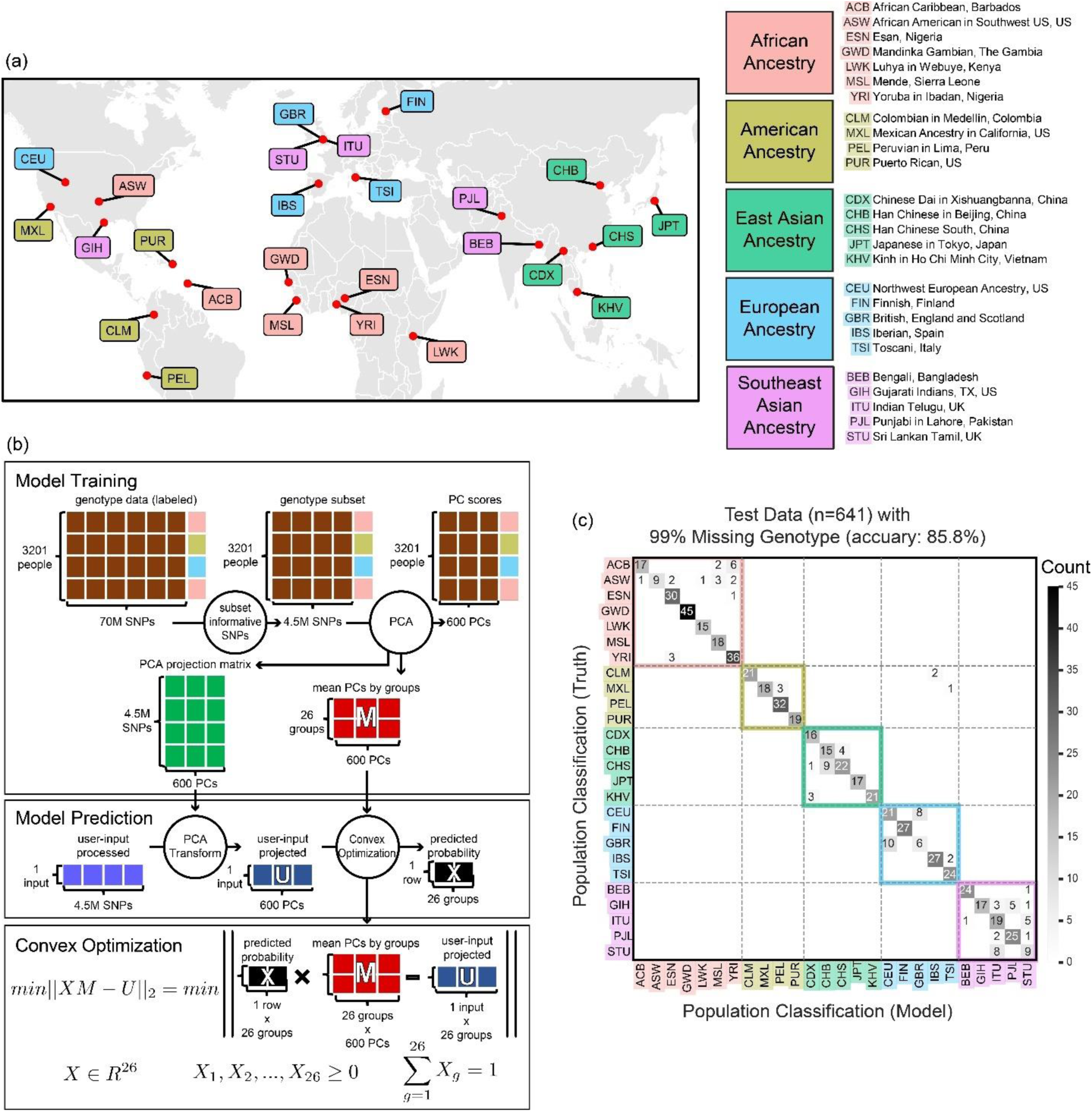
Schematic of model training and inference. (a) The geographic locations of the 26 populations in the 1KGP data set are shown in the truncated world map, with 5 color codes based on geographic regions. The right side shows the population description and three letter code that will be used throughout the paper. (b) Schematic of Model Training is shown on top. The training data is shown as a matrix of 3201 individuals by 70 million genotypes at SNP sites. We first subset the SNPs to keep 4.5 million ancestry-informative SNP sites as described in section 2.2 of Methods. We then conducted PCA to reduce the dimensionality to 600 principal components (PCs). Model prediction is shown on the bottom. An example of a user input of 1 sample is shown although users can input an arbitrary number of samples. The input data is first centered using the mean vector from the training data, imputed for sites with missing genotypes, and scaled appropriately as described in section 2.5 of Methods. Convex optimization is used to compute the contributions of the 26 population groups to the ancestry of the input data with the constraint that the contributions are non-negative and sum to 1. (c) The confusion matrix of classification results after splitting the 1KGP data into training-test (80-20%) is shown. Each test input is classified as the member of the population group with maximum contribution to that individual. Missing data was simulated by randomly removing 99% of the SNP sites from the data. Classification accuracy with 99% missing data is 86% (with no missing data, the accuracy is 89%), and all the misclassifications except for three cases are still within the same geographic region. Supplementary Figure 1 shows the confusion matrix generated with different degrees of missing genotypes.

In its final form, our method scAI-SNP (pronounced sky-snip) is remarkably easy to use. The user can take single-cell data of interest and extract the genotype of the 4.5 million ancestry-informative SNP sites. We genotype the SNP sites of scRNA-seq samples using a simple extension of the tool SComatic^24, 25^. Other methods such as Monopogen^25^, an elegant work that uses linkage disequilibrium to accurately detect germline variants, can also be used. Importantly, as shown below, the inference is remarkably robust to missing data and sequencing errors and has little diminished accuracy when as many as 99% of the SNP sites are not detected.

To carry out the inference of ancestry, we simply multiply the genotype vector of the 4.5 million SNP sites (allele vector) by a 4.5 million by 600 dimensional PCA projection matrix to obtain a 600-dimensional vector representation of the input data. We then use convex optimization to find the linear combination of the 26 mean vectors of each population group that best approximates the input vector with the constraint that the coefficients of the linear combination are non-negative and sum to 1. The computed coefficients are the contributions of the 26 population groups to the ancestry of the individual from whom the single-cell data was obtained. A similar approach for linear deconvolution of ancestry has been applied to frequencies of variants^26^ and principal components derived from these frequencies^27^ but at the level of populations as opposed to individuals. We will show that scAI-SNP works for different modalities of single-cell profiling methods, across different tissues and cell types, and even when applied to data from cancer cells. The code for scAI-SNP is available on the Hormoz Lab Gitlab page^28^ (https://gitlab.com/hormozlab/scAI-SNP).

## 2. Methods

scAI-SNP involves two stages: 1) model training and 2) model prediction. Briefly, for model training, we used the 1KGP dataset to generate a set of ancestry-informative SNPs and train an algorithm which takes in a vector corresponding to the alleles at these SNP locations and outputs a probability distribution over the 26 ancestral groups. For model prediction, we first used an extension of SComatic^24^ to genotype the ancestry-informative SNP sites in single-cell data, thereby obtaining the input vector to the model and then applied scAI-SNP to infer ancestry. Next, we describe each step of training and prediction in detail.

### 2.1 Data source for training

The 1000 Genomes Project (1KGP), launched in 2008, has collected and published whole-genome sequence (WGS) data to enhance our understanding of the genetic contribution to human phenotypes^23^. The New York Genome Center (NYGC) used 3202 samples from this project to conduct whole-genome sequencing at 30X coverage, followed by variant calling, and has made the data publicly available^29^. We excluded one individual who was reported as admixed of two population groups, reducing the number of individuals to 3201, each of whom was assigned to a single population group from the 26 different population groups used in the study (Figure 1a). From the 88 million genetic variations (SNPs, insertions and deletions, and structural variants) present in the data, we only used 70 million autosomal SNPs.

### 2.2 Data Pre-processing

To identify the ancestry-informative SNPs, we discarded SNPs where the most common genotype within a population was the same across all 26 population groups. This reduced the number of SNPs from approximately 70 million to 4.5 million; a set that we refer to as the ancestry-informative SNPs. Across the 3201 individuals, each of these SNPs was either the reference genotype (e.g. A) or a specific alteration (e.g. A to a C mutation). We did not see multiple alternative alleles in the dataset for a given SNP. Our procedure for reducing the number of SNPs may throw out SNPs that are unique to a particular group (and therefore informative about ancestry) but are not the most common genotype within in the population. Nevertheless, we enforced this rule to minimize computation without comprising the accuracy of our inference. At least 74 individuals were included in each population group (geographic distribution shown in Figure 1a and the numbers are listed in Supplementary Table 1).

As shown in Figure 1a, the 26 populations can also be grouped by 5 different geographic regions: Africa, America, East Asia, Europe, and Southeast Asia, where “region” is defined as the regional ancestry, not where the individuals reside. For example, several populations in the United Kingdom, such as Indian Telugu and Sri Lankan Tamil, would be from Southeast Asian ancestry, not European.

Next, we converted the genotypes of the 4.5 million ancestry-informative SNPs for each individual from a vector of base-pairs to a vector of numbers. For a given individual and a given SNP, if both base pairs (maternal and paternal alleles) matched that of the reference genome GRCh38, we assigned a 0. If only one of the base pairs (either from the maternal or paternal allele) matched the reference, we assigned a 0.5. Otherwise, when both did not match the reference, we assigned a 1. This numerical conversion does not discard any information about the mutations, as described above, we either see the reference or a single alternative allele for each SNP in the in the dataset. This is a reasonable numerical representation because the genotypes that an individual inherits are linear combinations of the genotypes of their parents.

Having obtained a numerical representation for the SNPs for every individual, we then centered the data. To do so, we thought of the data as a 3201-individual by 4.5-million SNP matrix. We computed the average of each column, corresponding to the average value of each SNP across the 3201 individuals. We then subtracted the average value of each column from all the entries of that column.

### 2.3 Inference algorithm

In addition to the description below, the model training and inference algorithm are depicted visually in Figure 1b.

#### 2.3.1 Principal component analysis (PCA)

We used Principal Component Analysis (PCA) to transforms the high-dimensional data (3201 individuals by 4.5 million features) into a lower-dimensional data of 3201 individuals by fewer features using an orthogonal transformation followed by truncating the feature space. This step is essential to reduce the computational burden when using the model to compute predictions for user-provided inputs. However, conducting PCA on 4.5 million features is challenging because, first, we have to compute a 4.5 million by 4.5 million mutation covariance matrix and then diagonalize that matrix. Instead, we computed the covariance across individuals which is a 3201 by 3201 matrix (referred to as the Gram matrix) and used this matrix to compute the principal components in the 4.5-million dimensional feature space.

The centered data matrix *X* is n by p, where n is the number of individuals (3201 in this case), and p is the number of SNPs sites (4.5 million in this case). We can use singular value decomposition to express the data matrix as:

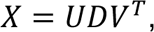

Where columns of *U* are the eigenvectors of the gram matrix (*XX^T^*), columns of *V* are the eigenvectors of the covariance matrix (*X^T^X*), and *D* is a *n* by p matrix in which the first (*n* – 1) diagonal entries are square-roots of the eigenvalues of *XX^T^* or *X^T^X* and all other entries are zero. Our goal is to compute *V*′, a truncated version of *V* with the first 600 eigenvectors of the mutation covariance matrix, to conduct PCA. Because directly computing *V* is computationally expensive, we first compute *U*, the eigenvectors of the 3201 by 3201 Gram matrix. We keep the first 600 eigenvectors with the largest eigenvalues to construct a subset of *U*, *U*′, as a 3201 by 600-dimensional matrix. *D*′ is a diagonal 600 by 600-dimensional matrix, where the diagonal entries are the first 600 largest eigenvalues. The choice of truncating to 600 dimensions is based on a sensitivity analysis shown in the Supplementary Table 2.

We can express *V*^′^ in terms of *U*^′^,

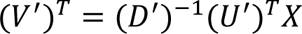

Where we have used the fact that (*U*′)*^T^U*′ = *I* and (*D*′)^−1^ is the inverse of matrix *D*′. It follows,

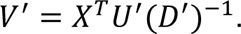

*V*′ can be used to project a vector in the original 4.5-million-dimensional feature space to a 600-dimensional space. We will use vectors in this lower-dimensional space to maps vectors of alleles to their ancestry.^31^

#### 2.3.2 Ancestry inference

To infer ancestry, our algorithm takes as input the data after PCA (600-dimensional input) and outputs the contributions of the 26 population groups to the ancestry of the input sample. This approach is appropriate because an individual’s ancestry can belong to more than one of the population groups.

The predicted ancestry is computed by solving the following constrained optimization problem using convex optimization:

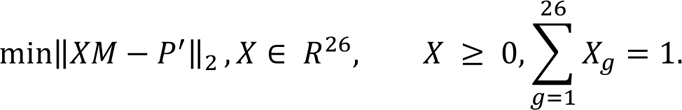

The parameters of interest, *X*, is a 26-dimensional non-negative row vector whose elements sum to 1, making it a vector of probabilities. *X_g_* is the contribution of population group *g* to the input. *M* is a 26 by 600 matrix in which row *i* is computed by projecting all the individuals in population group *i* in the training data after pre-processing into the PC space and averaging them. *M* then captures the average representation of each population group in the training data. *P*′ is a row vector of 600-dimensional user-input that has been pre-processed and projected into the PC space of the training data. Note that this optimization can be generalized for *X* to be a matrix of *n* row vectors, with *n* rows of user-input vectors (*P*′) in a form of row matrix in the scAI-SNP method. The optimization problem that we solve is finding the linear combination of the vectors representing each population group in the training data that most closely resembles the input vector with the coefficients constrained to be non-negative and summing to 1. We implemented convex optimization using package cvxpy (version 1.3.2) in Python 3.7.4 with a L2 norm to solve for vector *X*^32^. In cases where we aimed to classify the input to only one population group, we selected the population group *g* for the input with maximum *X_g_*.

### 2.4 Imputation

Because scRNA-seq data is expected to be sparse, we need to account for missing genotypes by imputation. To test our imputation approach, we simulated various degrees of missing data. We first split the data randomly into 80-20% proportion of training and test data, respectively, and implemented varying amounts of missing data in the test data (90%, 99%, and 99.9% of the allele calls for ancestry-informative SNPs missing). To implement missing data, for each test datapoint, we randomly selected a subset of SNP sites and marked them as missing. This subset can vary from one datapoint to another. We first imputed all the missing values of the SNPs with the corresponding genotype mean for each SNP site in the training data and then centered the data. This ensures that the missing SNP sites are non-informative for dimensionality reduction and classification. After PCA projection, we corrected for the missing data by multiplying the inverse of the proportion of the observed data (for example a factor of 100, for 99% missing) to scale the principal components. Supplementary Figure 2 shows the first four principal components for the original test data and the imputed and corrected test data. Even with 99% of data missing at random, the principal components in the first four dimensions are quite similar to those of test data. We computed the downstream effects of missing data on the inference accuracy and demonstrate them in Figure 1c and Supplementary Figure 1.

### 2.5 Number of Principal Components

To determine the ideal number of principal components to use, we conducted a sensitivity analysis in which we split the 3201 individuals into training and test sets at random (80-20% split) and conducted dimensionality reduction and classification with varying numbers of principal components. As shown in Supplementary Table 2, using the first 600 principal components is sufficient as the inference accuracy does not increase significantly with additional principal components.

### 2.6 Genotyping

Users may use their own genotyping algorithm to prepare inputs for scAI-SNP. The list of 4.5 million SNP sites and their information is provided as a text and Browser Extensible Data (BED) file in the scAI-SNP GitLab page^28^. To genotype SNP sites in single-cell data sets, we extended SComatic. Specifically, users can directly use SComatic to readily prepare their own single-cell data for scAI-SNP. SComatic conducts variant calling of the given 4.5 million sites with various requirements to confidently assign a genotype for each site. It requires 5 reads at minimum for proper genotype extraction, and if the variant base calls to read depth ratio is less than 0.1, the site is considered to be homozygous reference (0/0). Similarly, if the ratio is greater than or equal to 0.1 but less than 0.9, the SNP is considered to be heterozygous mutation (0/1) and homozygous mutation (1/1) if the ratio is greater than or equal to 0.9.

### 2.7 Data sources for testing

#### 2.7.1 Bone marrow data set

We obtained bone marrow aspirates from 5 healthy individuals (BM_ID1 to BM_ID5) of diverse backgrounds. These healthy donor samples were purchased as de-identified samples from the Boston Children’s Hospital. Mono-nuclear cells were isolated from each sample and processed as previously described in^30^. In addition to these 5 samples, we used scRNA-seq data and WGS profiling of bone-marrow cells from a previous study on JAK2-mutant myeloproliferative neoplasms (MPN)^30^ as samples BM_ID6 and BM_ID7. The WGS had an average depth of 32X. For BM_ID4 to ID7, CD34 positive cells were enriched from the mono-nuclear cells and used to generate the single-cell libraries. To call the alleles for the ancestry-informative SNPs in the WGS data we used Strelka2 and set a minimum genotype quality (Phred score) of 20 or higher with a depth of 3 reads or higher.

#### 2.7.2 Heart single-cell dataset

A study of cardiovascular disease that sought to characterize genetic heterogeneity of heart cells conducted single-cell RNA sequencing of 14 donors with unremarkable cardiovascular disease history^31^. CD45 positive cells were enriched from various regions of the heart, processed using the 10X Genomics platform, and sequenced using HiSeq 4000 (Illumina) and NextSeq 500 (Illumina) with a minimum depth of 20,000 to 30,000 read pairs per cell. Each sample had about 31,600 cells on average. Our study used 76 samples that had at least 20,000 ancestry-informative SNPs detected, which were sampled from six different regions of the heart.

#### 2.7.3 GTEx dataset

As part of the Genotype-Tissue Expression (GTEx) project, a study produced snRNA-seq profiles of eight tissue types from 16 donors^32^. Each of these snRNA-seq samples has about 13,700 nuclei with 1,500 reads per nucleus on average. On average about 1.35% of the ancestry-informative SNP sites were detected in the data. We used 10 samples from 4 donors that had at least 20,000 detected sites.

#### 2.7.4 Ovarian cancer MSK SPECTRUM dataset

We analyzed scRNA-seq data from 160 multi-site primary and metastasized tumor tissues originated from 42 patients with high-grade serous ovarian cancer (HGSOC)^33^. This dataset also had matched WGS data for normal peripheral blood mononucleated cells and tumor cells. Chromium Single-Cell 3’ Reagent kit v3 (10X Genomics) was used to sequence the data. We used 268 samples from 26 sites and 41 patients in total. 6 samples that had fewer than 20,000 ancestry-informative SNP sites detected were discarded.

#### 2.7.5 Pan-tissue scATAC-seq dataset

The pan tissue scATAC-seq data was generated from a study that created a cell atlas of *cis*-regulatory elements^34^. The study team conducted sci-ATAC-seq, a modified scATAC-seq that uses combinatorial indexing, with 92 samples from 30 tissue types^35^. We used 66 samples from 4 donors that were processed and used in the SComatic study^36^. As with the other data sets, low-quality scATAC-seq samples, which had fewer than 20,000 of the 4.5-million ancestry-informative SNP sites detected, were discarded (47 samples out of 66 representing 22 tissue types passed this quality control).

## 3. Results

We trained scAI-SNP on the 1KGP data as described in Methods. Briefly, the data contains 70 million SNPs across 3201 individuals who belong to one of 26 population groups (Figure 1a). We identified a subset of 4.5 million SNP sites that were informative for ancestry inference (the most common allele within each population group was not the same across all the groups) and captured each individual’s genetic information into a vector of alleles of these 4.5 million SNP sites. We then used principal component analysis (PCA) to reduce the dimensionality of the allele space from 4.5 million to 600. To make an inference, the user provides an input vector corresponding to the alleles of the 4.5 million SNP sites for a given sample. This input can contain missing values if some of the SNP sites were not genotyped. Our algorithm replaces the values of the missing SNP sites with the mean values from the training set. The prediction is computed by multiplying this input vector by a 4.5 million by 600-dimensional matrix to obtain the projection of the input vector to the 600-dimensional PCA space. We appropriately scale the resulting principal components to account for the missing data by multiplying by the inverse of the proportion of the observed data (for example a factor of 100, for 99% missing). We then use convex optimization to find the linear combination of the mean vectors of each population group that best approximates the input vector. We constrain the coefficients of this linear combination to be non-negative and sum to 1. The resulting 26-dimensional output corresponds to contribution of each of the 26 population groups to the donor from whom the sample was obtained (Figure 1b).

To validate scAI-SNP, we first generated synthetic test data from the 1KGP dataset itself. We split the dataset into a training-test set (80-20% split) and used the training set to train the model. We implemented missing alleles in the test data by dropping the value of 90%, 99%, and 99.9% of randomly chosen SNP sites for each test data point. With no missing data, the accuracy was 89%, which dropped to 86% with 99% of the data missing (Supplementary Figure 1 and Figure 1c). Therefore, the predictions are remarkably robust to missing data. Importantly, the incorrect predictions of the model are still consistent with expected human demographics and migration patterns. For example, as shown in Figure 1c, individuals from the ASW (African American in the Southwest of US) group are sometimes mistakenly inferred to be from the ESN (Esan, Nigeria) and the YRI (Yoruba in Ibada, Nigeria) groups. Similarly, individuals from the GBR (British, England and Scotland) group are mostly attributed to CEU (Northwest European Ancestry, US). Similarly, CHS (Han Chinese South) population and CHB (Han Chinsese in Beijing) populations also demonstrate similarities and mixtures. From known migration histories of these populations, such predictions errors are not necessarily biological and can potentially be attributed to admixtures and limitations in the self-reported ancestry in the 1KGP data. We reasoned that a more complex model with higher accuracy will most likely overfit the 1KGP dataset and not generalize as well. Additionally, our linear model correctly infers the ancestry of mixed individuals (Supplementary Figure 4). Therefore, we proceeded with our simple linear fit for ancestry inference. An alternative approach would have been to remove the recently admixed populations (such as ASW) from the inference or to predict ancestry up to the 5 geographic regions. We decided against this approach because, as we discuss in more detail in the Discussion, the predicted ancestry can also be used as a proxy for other contexts for an individual, such as environment and socioeconomic status. Therefore, we decided to predict the contributions of all 26 populations groups in the 1KGP dataset including the recently admixed population groups.

To determine the limitation of our method, namely approximating the ancestry of an individual as a linear combination of the mean of the 26 population groups in the 1KGP dataset, we first set out to characterize the heterogeneity in ancestry of individuals within the same population group. To do so, we plotted the first two principal components of the 3201 individuals in the 1KGP dataset computed above. As shown in Figure 2a, the PCA plot shows distinct groups that correspond primarily to the 5 geographic regions sampled. However, as previously reported^11^, we observe heterogeneity within each population group and that many populations form a continuum of points as opposed to distinct clusters. To quantify the degree of heterogeneity in each population group, we split the 1KGP dataset to a training and test data sets (80-20% split) and computed the principal components only using the training data. We then computed the similarity of the test points to the mean of their population group by computing the cosine similarity between the vector corresponding to test point in the PC space and the mean of all vectors in the training data in the same population group as the test point (Supplementary Figure 3). A value close to 1 indicates a high degree of similarity whereas a value close to 0 indicates low similarity. Certain population groups, particularly those with American and Southeast Asian ancestry, exhibited high heterogeneity in that many individuals in those population groups had low similarity to the population mean.

**Figure 2.**
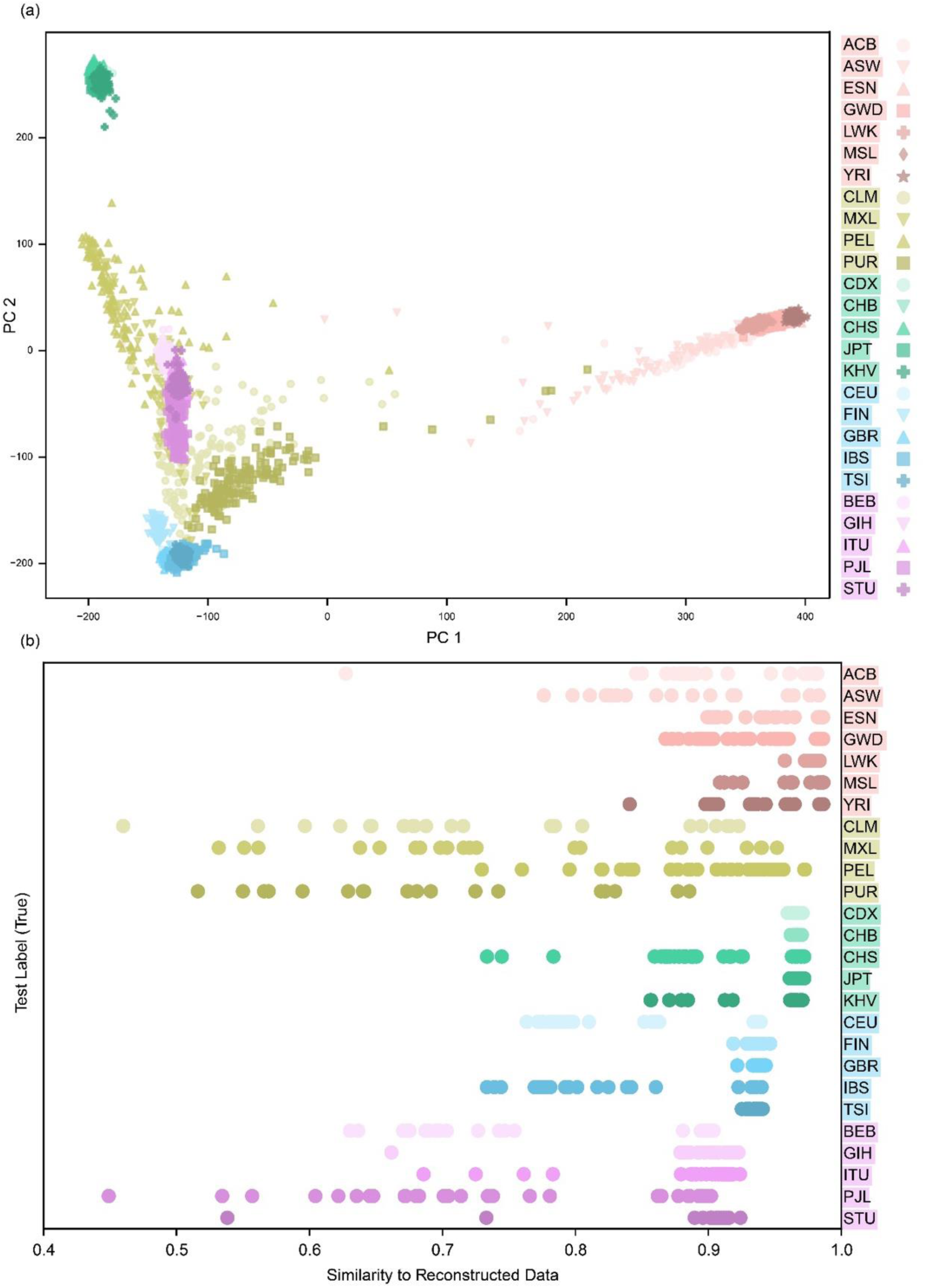
PCA of the training data and the heterogeneity within each population group. (a) The first two principal components of the 1KGP data. The marker shape and colors capture the 26 different population groups, with a different color used for each of the 5 different geographic regions. The PCA plot shows a clear separation of the points by the 5 different regions. However, the PCA plot also shows the admixed nature of the populations and that many of the population groups form a continuum of points as opposed to distinct clusters. (b) After splitting the data into training-test (80-20%), we converted each test data point to a vector in the 600-dimensional PCA space obtained from the training data. We used convex optimization then to construct the closet possible vector to the test vector using a linear combination of the mean vectors of the 26 population groups, where the coefficients of the linear combination were constrained to be non-negative and sum to 1. For each test data point, we have plotted the cosine similarity of the vector corresponding to the test data point and its reconstruction. The values range from 0 to 1, where 1 indicates that the data can be perfectly explained by the mean PC vectors of the 26 population groups. Lower values of similarity result from heterogeneity of the data within each population group and lack of population groups in the training data that would explain this heterogeneity through admixture.

We reasoned that some of this heterogeneity could be explained by admixtures between the 26 population groups present in the training data. To remove this contribution from the heterogeneity measure, we used scAI-SNP to obtain the closest approximation of each test point as a linear combination of the mean vectors of the 26 population groups. Figure 2b shows the cosine similarity between each test vector and its best approximation. Although the level of heterogeneity decreased, individuals from American and Southeast Asian ancestry are still poorly approximated as linear combinations of the mean vectors of the 26 population groups, most likely because these groups are admixtures with populations groups other than the 26 used in the training data. Therefore, the inferred ancestry labels for these population groups (such as CLM, Colombian in Medellin, Colombia, and PJL, Punjabi in Lahore, Pakistan) should be interpreted with caution because of the high degree of heterogeneity in the training data. We propose ways to supplement the 1KGP dataset in the Discussion section. Nevertheless, for many applications our algorithm is sufficiently accurate especially when inferring up to the 5 primary geographic regions.

To test the accuracy scAI-SNP on single-cell data and on individuals outside of the training data, we collected bone-marrow biopsies on 7 individuals with self-reported race (3 White, 1 Black, 2 Asians, and 1 Indian; Methods). We performed single-cell RNA sequencing of bone marrow mono-nuclear cells (MNCs) from all these samples and applied scAI-SNP to the data. As shown in Figure 3a, the inferred ancestry of each sample is consistent with the self-reported race. To determine whether scAI-SNP predictions from sparse single-cell data is comparable with predictions from whole genome sequencing, we obtained WGS data from peripheral blood of 2 of the 7 individuals. The 4.5 million SNP sites in the WGS data was genotyped using Strelka2 (version 2.9.10)^37^. We obtained allele calls for 99.3% and 97.6% of the 4.5 million ancestry-informative SNP sites of the 2 donors from WGS data compared with 13.0% and 7.43% obtained from their single-cell data respectively. As shown in Figure 3b, the inferred ancestry of these two individuals from single-cell RNAseq data agree with that inferred from WGS data despite the sparsity of the single-cell data. Taken together, scAI-SNP provides accurate inference of ancestry from single-cell RNAseq data that is consistent to that inferred from WGS data.

**Figure 3.**
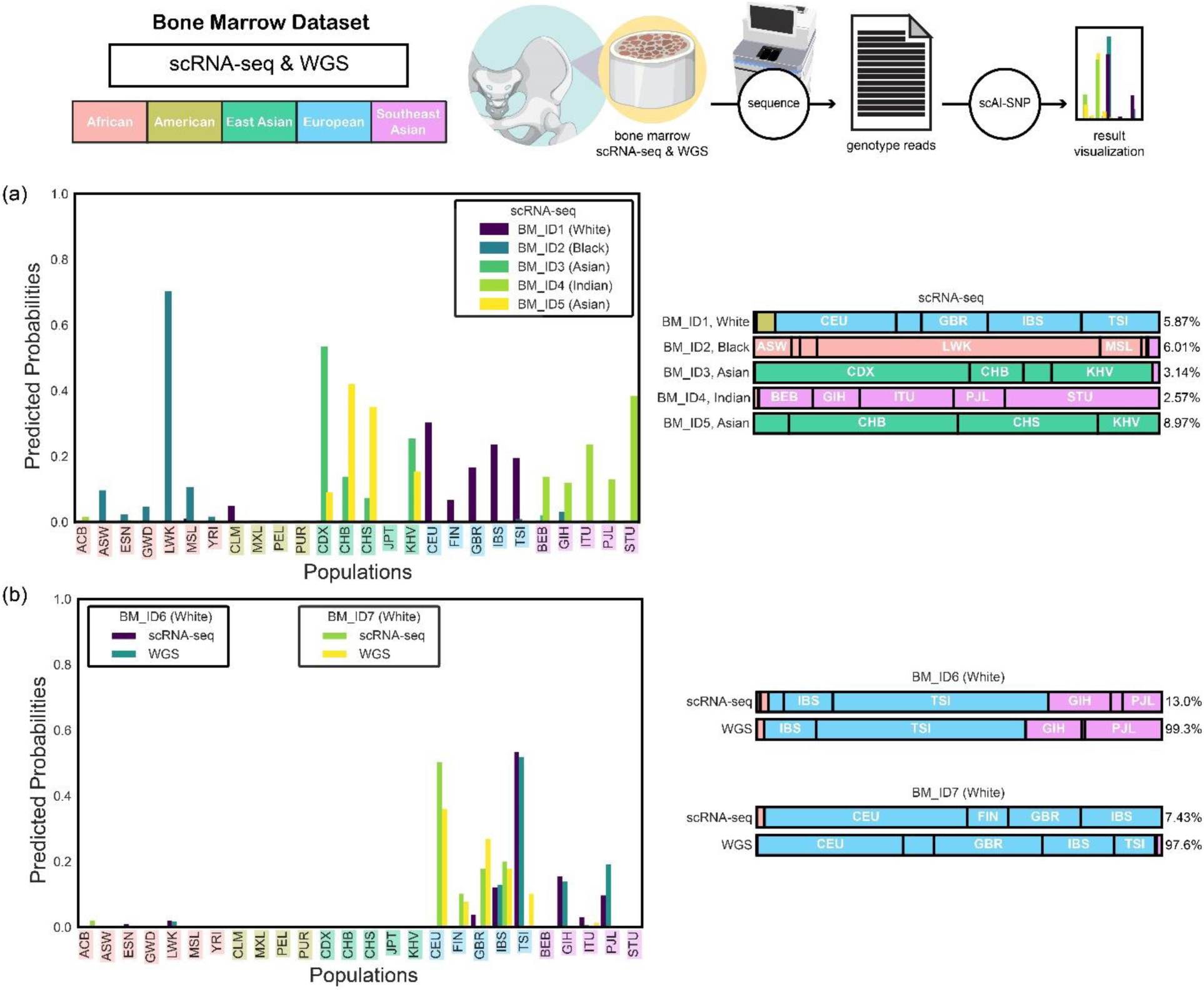
Validation of scAI-SNP for ancestry inference. (a) Inferred ancestry from single-cell RNAseq data of bone marrow mononuclear cells from 5 donors with known self-reported race. The vertical bar plots depict the predicted contribution of each of the 26 population groups to each donor. These probabilities are plotted again in horizontally stacked bar plots on the right in a more succinct form. The percentages on the right side are the fraction of the 4.5 million ancestry-informative SNP sites detected in each sample. (b) Same plots as in (a) for 2 samples from donors who self-reported as White. Ancestry inferred from whole genome sequencing of DNA extracted from the peripheral blood of these donors yielded results consistent with those from scRNA-seq samples.

Next, we set out to determine whether scAI-SNP can infer ancestry from scRNA-seq data from a wide variety of cell types and tissues. To do so, we applied scAI-SNP to scRNA-seq data from the Heart Cell Atlas dataset^31^. We used data from 14 individuals who had between 3 and 6 scRNA-seq samples (78 in total) from cells sampled from different regions of the heart (Figure 4 and Methods). These samples had on average ∼31,600 cells and were sequenced with a minimum depth of 20,000 to 30,000 read pairs per cells. We detected on average 2.1% of the 4.5 million SNP sites in these samples. 2 samples that had fewer than 20,000 sites (or 0.44% site detected) were not used for further analysis. Figure 4a shows the predicted ancestry for each donor across all their samples as a pie-chart. The inferred ancestry was consistent across different samples from the same individual. The fact that 13 out of the 14 individuals in the Heart Cell Atlas Dataset is of European ancestry highlights the need for inferring ancestry of single-cell data to ensure diversity of atlases. To further validate this finding across more diverse tissues, we inferred ancestry of individuals in the GTEx dataset who had multiple single-nucleus RNA sequencing (snRNA-seq) data obtained from different tissues^32^. Each snRNA-seq sample had about 13,700 nuclei with 1,517 reads per nucleus on average, and we detected about 1.35% of the SNP sites and discarded the samples that didn’t have at least 20,000 ancestry-informative SNP sites detected. We applied scAI-SNP to data from individuals that had samples from multiples tissues (4 individuals with 10 samples) as shown in Figure 4b. Again, we observed consistent inferred ancestries across different tissues from the same individual. Taken together, scAI-SNP can consistently infer ancestry from scRNA-seq and snRNA-seq data from cells from a wide variety of tissues.

**Figure 4.**
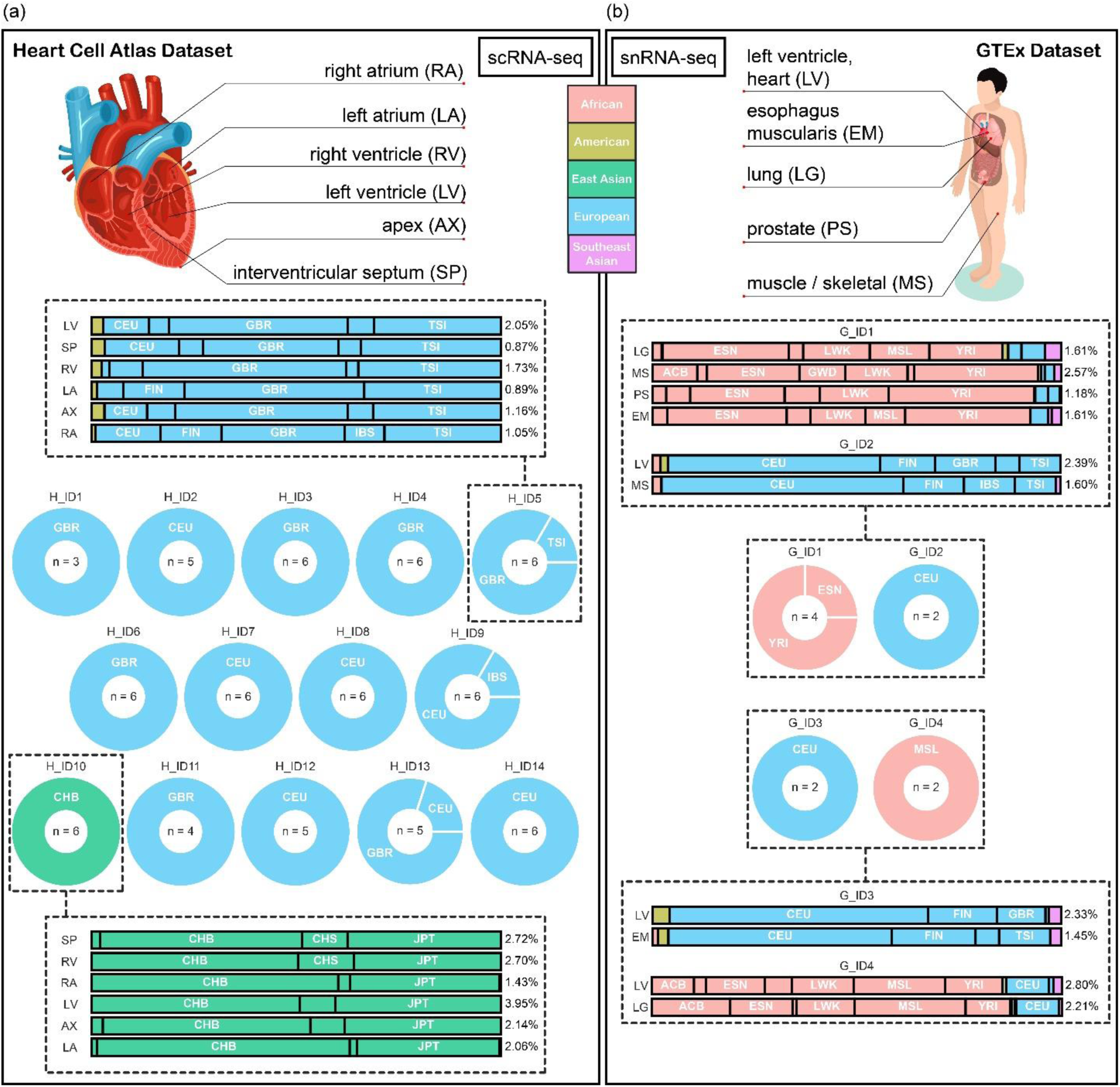
scAI-SNP results from single-cell RNA sequencing of a diverse set of cell types and tissues. (a) The inferred ancestry of 14 individuals (H_ID1 to H_ID14) from their scRNA-seq data in the Heart Cell Atlas dataset. Each individual had multiple samples obtained from different regions of the heart as shown approximately in the schematic. The pie charts show the distribution of classifications (population group with the highest predicted probability) across all the samples for each individual. The number of samples for each individual is shown in the center of the pie chart. For individuals H_ID5 and H_ID10, the detailed predictions are shown as the horizontally stacked probability plots, where the left labels indicate which region of the heart was sampled and the right shows the fraction of ancestry-informative SNP sites that were detected. (b) The inferred ancestry of snRNA-seq data from the GTEx dataset. The labels on the left side of the horizontal bar plots show the region from which the sample was obtained with the legend included with the simplified human schematic. The schematic is an approximation and does not show the exact regions from which the samples were obtained.

To determine whether we could accurately infer ancestry from scRNA-seq data of cancer cells, we analyzed the MSK SPECTRUM Ovarian Cancer dataset^33^. From this dataset, we used data from 41 individuals who had had 3 to 12 scRNA-seq samples and corresponding WGS data (from both tumor and normal cells). The scRNA-seq samples contained about 17,100 cells on average and had about 2.0% of the ancestry-informative SNP sites detected. As before, 6 samples that had fewer than 20,000 ancestry-informative SNP sites detected were discarded. As shown in Figure 5, all 41 individuals had consistent ancestry inferred up to the 5 geographic regions. For 39 out of the 41 individuals, WGS data of both tumor and normal cells were available. Among these 39 individuals, 36 individuals’ tumor and normal cell ancestry were consistent (the population group with the highest inferred probability was the same). The predicted ancestry of the three individuals that were inconsistent between tumor and normal cells were still within the same geographic region. Similarly, in the cases where the most probable inferred ancestry group from single-cell sample did not match that of the WGS normal data, the most probable groups were still within the same geographic region. Therefore, scAI-SNP can accurately and consistently infer ancestry from both healthy and tumor samples.

**Figure 5.**
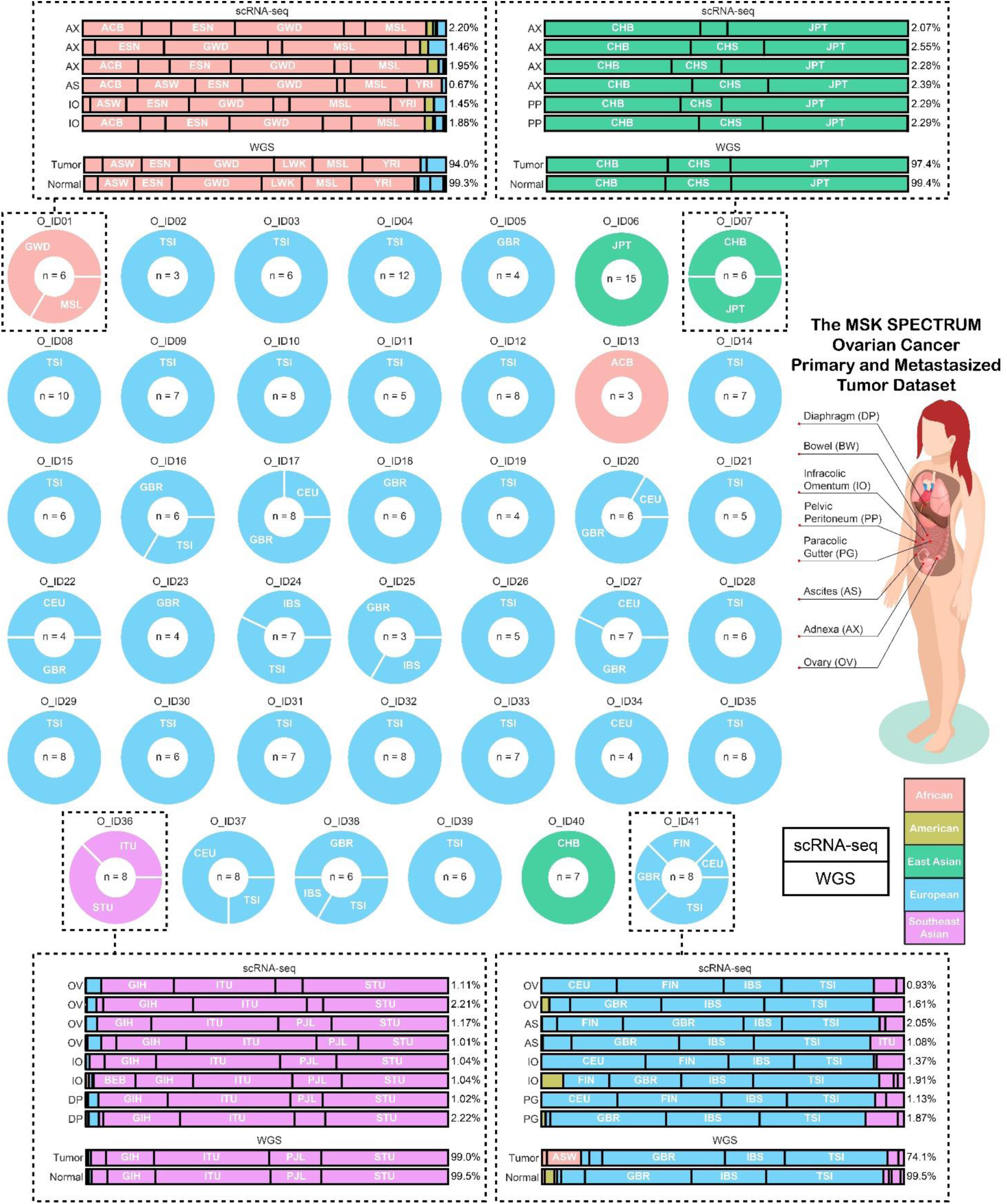
scAI-SNP results from scRNA-seq and WGS of cancer cells. Pie charts show the inferred ancestry (ancestry group with the largest predicted probability) across all the scRNA-seq tumor samples for a given individual across 41 individuals in the MSK SPECTRUM Ovarian Cancer dataset. The number of samples for each individual is shown in the center of the pie chart. The schematic shows approximately the different regions from which the samples were obtained. For individuals O_ID01, O_ID07, O_ID36, and O_ID41, the stacked predicted probabilities are shown along with the labels on the left that indicate which tissue was used and on the right that show the fraction of the 4.5 million ancestry-informative SNP sites detected in each sample. Note that multiple scRNA-seq samples may originate from the same site. For those 4 individuals, we also show the inferred ancestry using their respective WGS of normal and tumor tissues.

Finally, we set out to determine if scAI-SNP could be applied to scATAC-seq data. To do so, we applied our algorithm to scATAC-seq data derived from cells collected from various types of tissues from 4 donors in the Pan-Tissue scATAC-seq dataset^34^. The samples contained on average roughly 114,800 cells and were sequenced to a depth of 6,500 raw reads per nucleus on average. As before, low-quality scATAC-seq samples, which had fewer than 20,000 of the 4.5 million SNP sites detected, were discarded. As shown in Figure 6, the inferred ancestry using scAI-SNP was consistent across all tissues for each of the donors. The inferred ancestries, up to the 5 major geographic regions, were consistent across all tissue types across all samples, as was the contribution of each of the 26 population groups to each sample qualitatively. Taken together with the results seen with other datasets, scAI-SNP can be applied to different modalities of the single-cell data, including scATAC-seq data.

**Figure 6.**
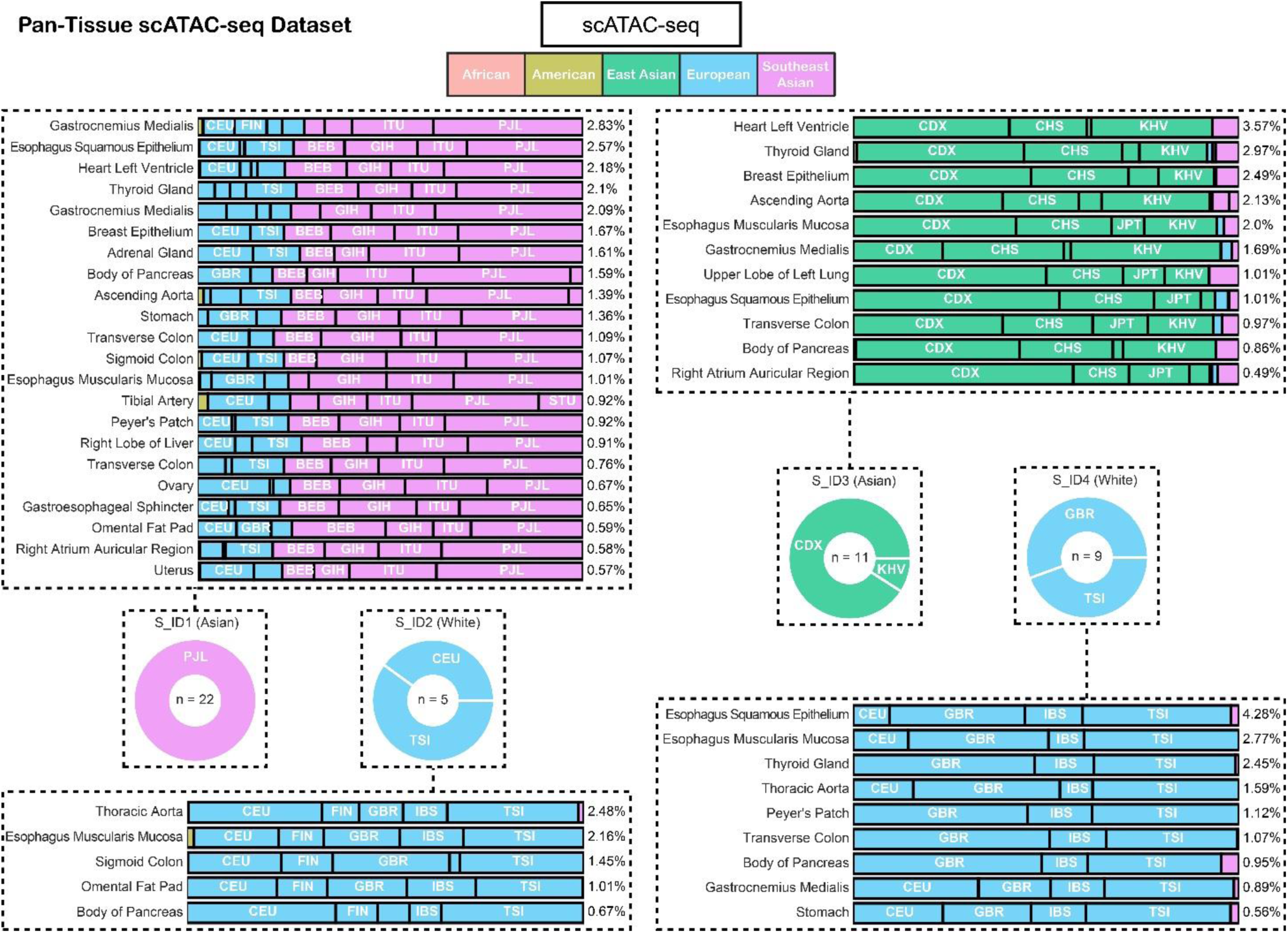
scAI-SNP consistently infers ancestry from scATAC-seq data from various types of tissues. Inferred ancestries of 4 donors’ scATAC-seq samples from various tissues are shown in the horizontal stacked plots. On the left side of the stacked bars are the description of tissues from which samples were collected and on the right side are the fraction of the 4.5 million ancestry-informative SNP sites detected in each sample. Each donor’s inferred ancestry across all of their samples is also summarized in the pie charts in the middle.

## 4. Discussion

Inferring ancestry from single-cell data is essential for constructing single-cell atlases that reflect human diversity. Here, we introduced a tool, called scAI-SNP, that takes in single-cell genomic data as input, extracts alleles from 4.5 million ancestry-informative SNP sites, and outputs a distribution over 26 ancestry groups, which represents the inferred contribution of each ancestry group to the donor of the sample. We showed that scAI-SNP is accurate and robust to the sparsity present in single-cell data, and can be applied to samples from various cell and tissues and even cancer cells sequenced using WGS, scRNA-seq, snRNA-seq, and scATAC-seq.

The key feature of scAI-SNP is the simplicity in its ancestry inference. We used the data from 3201 individuals in the 1000 Genomes Project (1KGP) to identify 4.5 million ancestry-informative SNP sites to learn linear transformations followed by convex optimization that converts an input allele vector of 4.5 million dimensions to a vector of probabilities over the 26 ancestry groups. This simple approach is justified because population admixtures (e.g. individuals with ancestors from different population groups) can be thought of as linear combinations of their parental groups. In addition, this simple approach minimized the risk of over-fitting to the training data. This is especially important because the ground truth in 1KGP comes from self-reported ancestries and is therefore limited in its accuracy.

Our approach has certain limitations. First, our method ignores any linkage disequilibrium between the ancestry-informative SNP sites. Accounting for linkage disequilibrium can significantly improve the ability to distinguish a true mutation from a sequencing error in single-cell data as shown recently by Dou et al.^25^, where the authors use their calls for germline mutations to validate global ancestry inference using PCA and carry out local ancestry inference using RFMix^18^. Although linkage disequilibrium is very informative if the goal is to call specific mutations with high confidence, for example for genetic association studies, we reasoned that for global ancestry inference the redundant information provided by the millions of ancestry SNP sites allow us to safely ignore linkage disequilibrium and use a much simpler approach.

Another important limitation of our work is that we cannot distinguish uncertainty in model prediction from admixtures of population groups. This is because the output of the model is a probability distribution over the 26 population groups in the training data (1KGP). A completely uninformative prediction would correspond to a uniform distribution over all 26 ancestry groups. An intermediate prediction of a distribution with significant weight on a subset of the ancestry groups can be interpreted both as model uncertainty across those groups or an individual who is an admixture of those groups. This issue is especially pronounced when the individual is from an ancestry group that is not one of the 26 population groups in the training data. Therefore, the biggest limitation of our method is the limited nature of our training data. 1kGPdata has been analyzed for several studies of ancestry, but its 26 populations do not represent an ideal sample of human diversity around the globe because they do not include the full range of genetic diversity. Our analysis of test data points cosine similarity to the closest linear combination of the mean vector of the 26 population groups highlights this. The heterogeneity in many population groups is consistent with missing population groups and that labels for many population groups (such as CLM, Colombian in Medellin, Colombia, and PJL, Punjabi in Lahore, Pakistan) can be misleading by neglecting the significant heterogeneity within those populations. In future iterations of scAI-SNP, we hope include more population from other studies of human genetic variation^38^, and potentially even data from All of US and UK Biobank^27^. Nevertheless, our simple tool can provide invaluable insight into ancestry information of single-cell datasets and guide the generation of single-cell atlases to represent human genetic diversity.

Finally, we outline broadly the benefits of ensuring diverse ancestry when constructing single-cell atlases. As described in this paper, it is rather easy to identify genetic variants that are predictive of an individual’s ancestry. However, this does not imply that traits, in particular neutral traits, can also be predicted from ancestry^39^. Intuitively, when we train models that predict ancestry from millions of genetic variants (i.e., the ancestry-informative SNPs defined in this work) we are accumulating small contributions to ancestry of each variant across all the variants. The contribution from each variant is positive and can be summed together without cancellation. Although a single variant is not predictive, many of them together can make accurate predictions. A trait is not like ancestry. The contribution of each variant to a trait can be both positive and negative. Therefore, accumulating the contributions of many variants to a trait is no more accurate than predicting the trait from a single variant. Naively, this argument implies that although ancestry can be inferred from genetic variants, ancestry itself is not useful for capturing diversity of single-cell phenotypes.

There are two reasons why this naive argument is not correct. First, rare variants that contribute significantly to a particular trait and are present predominantly in specific population groups due to hard sweeps such as selection or drift do occur and are clinically important. Examples are polymorphisms associated with Lactase persistence^40^, Duffy allele^41^ and G6PD deficiency that reduce the risk of malaria infection^42^, and EPAS1 gene in Tibetans for adaptation to high altitude^43^. In the clinic, some variants are predictive of propensity for disease, such as the 8q24 variant, which is associated with increased risk of prostate cancer and more prevalent in African Americans compared with individuals of European ancestry^44^. Such variation across population groups may also result in incompatibility of reference ranges computed using one population group when applied to another. For example, blood counts of individuals of European descent are significantly different compared with those of African descent^7^. Taken together, single-cell atlases need to incorporate and account for rare genetic variants across population groups.

Second, ensuring diversity of ancestry can indirectly ensure diversity of environment and socioeconomic status of the donors of atlases. For example, consider polygenic scores that aggregate the contributions of many genetic variants to a trait. The accuracy of polygenic risk scores varies significantly with context, such as age and income, in addition to ancestry^45^. In the cases where context is not recorded, ancestry can act as a proxy for environmental and socioeconomic factors. This would allow us, for example, to study impact of environment on biological traits, elucidating the interactions between context and genetic variation. Diversity also allows us to remove noise from linkage disequilibrium when associating variants with traits. This is highlighted by the fact that accuracy of polygenic scores decreases when applied to individuals with increasing genetic distance to the training cohort^46^. Ideally ancestry should be supplemented with race and ethnicity. Race and ethnicity are even more closely correlated with environment and socioeconomic status than ancestry, and with other social determinants of health such as racism and discrimination^47^. Constructing single-cell atlases from donors with diverse ancestry, alongside race and ethnicity, should ensure that the scientific discoveries made using such as atlases and their eventual use in the clinic result in improved and equitable health outcomes.

## Acknowledgements

We thank Sasha Gusev for helpful and insightful feedback on the manuscript. This work was supported by funding from the National Institutes of Health (NIH) National Heart, Lung, and Blood Institute grant no. R01HL158269, the Chan Zuckerberg Foundation (Seed Grant) CZF2019-002433, and the Chan Zuckerberg Initiative (CZI) Seed Networks Supplemental Funding Opportunity (Tissue Diversity Supplement) 2020-224559. F.M. and I.C.-C. thank EMBL for funding.

## Data availability

The code and scripts used for ancestry inference can be found on our lab’s GitLab page (https://gitlab.com/hormozlab/scAI-SNP).

The variants of 30x WGS data from the 1000 Genomes Project are publicly available at this link (https://www.internationalgenome.org/data-portal/data-collection/30x-grch38).

The scRNA-seq and whole-genome sequencing data used for the validation shown in Figure 3 (BM_ID6 and BM_ID7, also referred to as ET1 and ET2) is available via dbGAP under study accession number phs002308.

The scRNA-seq data of the Heart Cell Atlas is available on ENA (European Nucleotide Archive) with study accession number ERP123138 (https://www.ebi.ac.uk/ena/browser/view/ERP123138).

The sequencing data of GTEx samples is available on AnVIL (Analysis Visualization and Informatics Lab-space) and requires a data access application via dbGap under study accession number phs000424 (https://anvil.terra.bio/#workspaces/anvil-datastorage/AnVIL_GTEx_V9_hg38).

The WGS and scRNA-seq of ovarian cancer data set (MSK SPECTRUM) are available at this link (https://www.synapse.org/#!Synapse:syn25569736/wiki/612269).

The scATAC-seq data is available in the GEO database under accession number GSE184462.

The reference genome used (GRCh38) is available at this link. (https://hgdownload.soe.ucsc.edu/downloads.html).

---

**Supplementary Figure 1.**
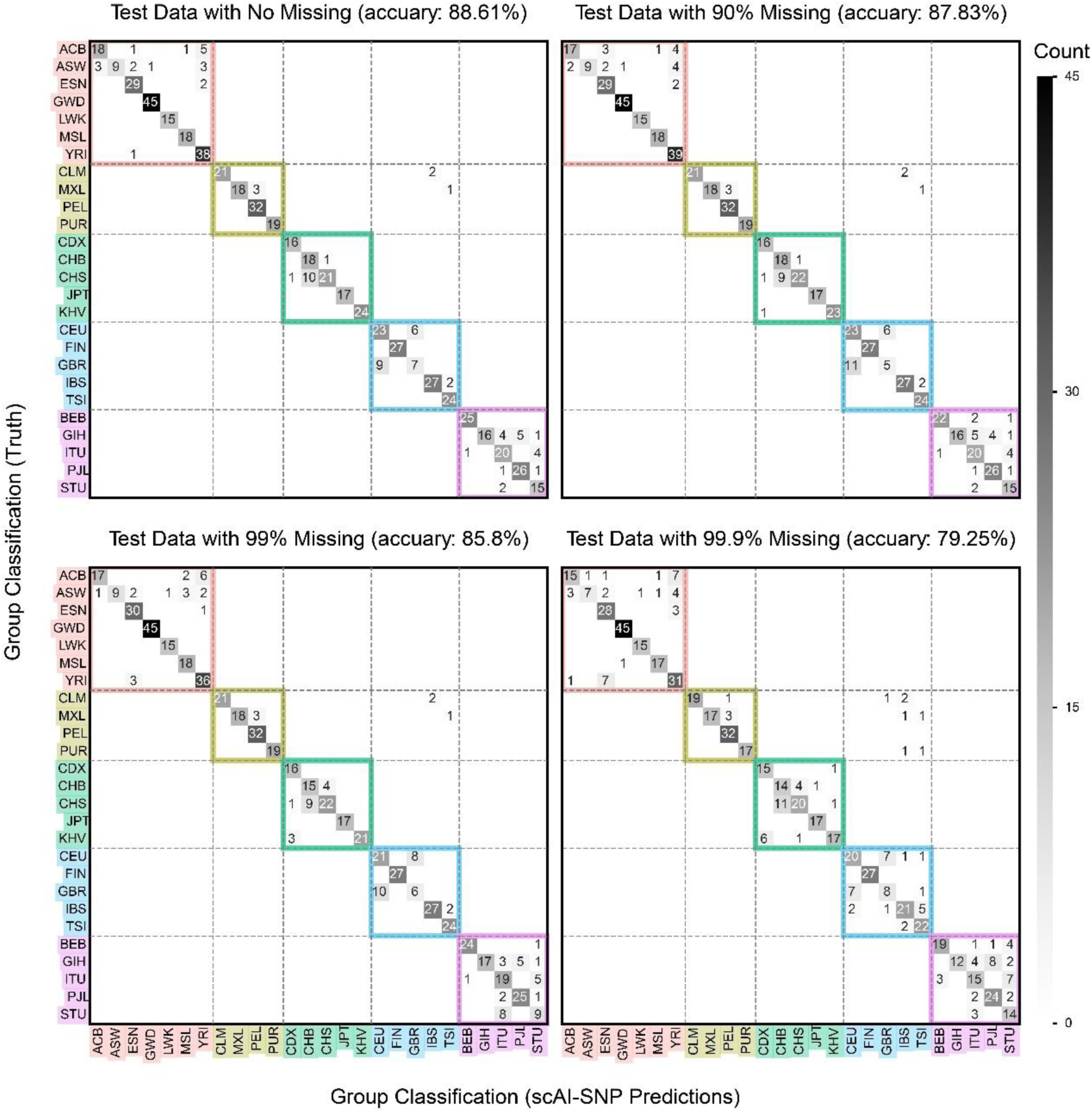
The confusion matrix of scAI-SNP ancestry inference results against the ground truth after training the model with 80% of the data and using 20% of the data as test data. In each confusion matrix, different degrees of missing data were used. Classification accuracies dropped from 88.6% to 79.3% from no missing data to 99.9% missing. The colors used here are the same with color coding in Figure 1 to 6. Off-diagonal entries inside the colored squares indicate intra-region misclassification, and off-diagonal entries outside the colored squares are inter-region misclassifications, which are rare. Most frequent misclassifications are in between GBR and CEU (British in the UK and European Americans in the US) and between CHS and CHB (Han Chinese South, China, and Han Chinese in Beijing, China).

**Supplementary Figure 2.**
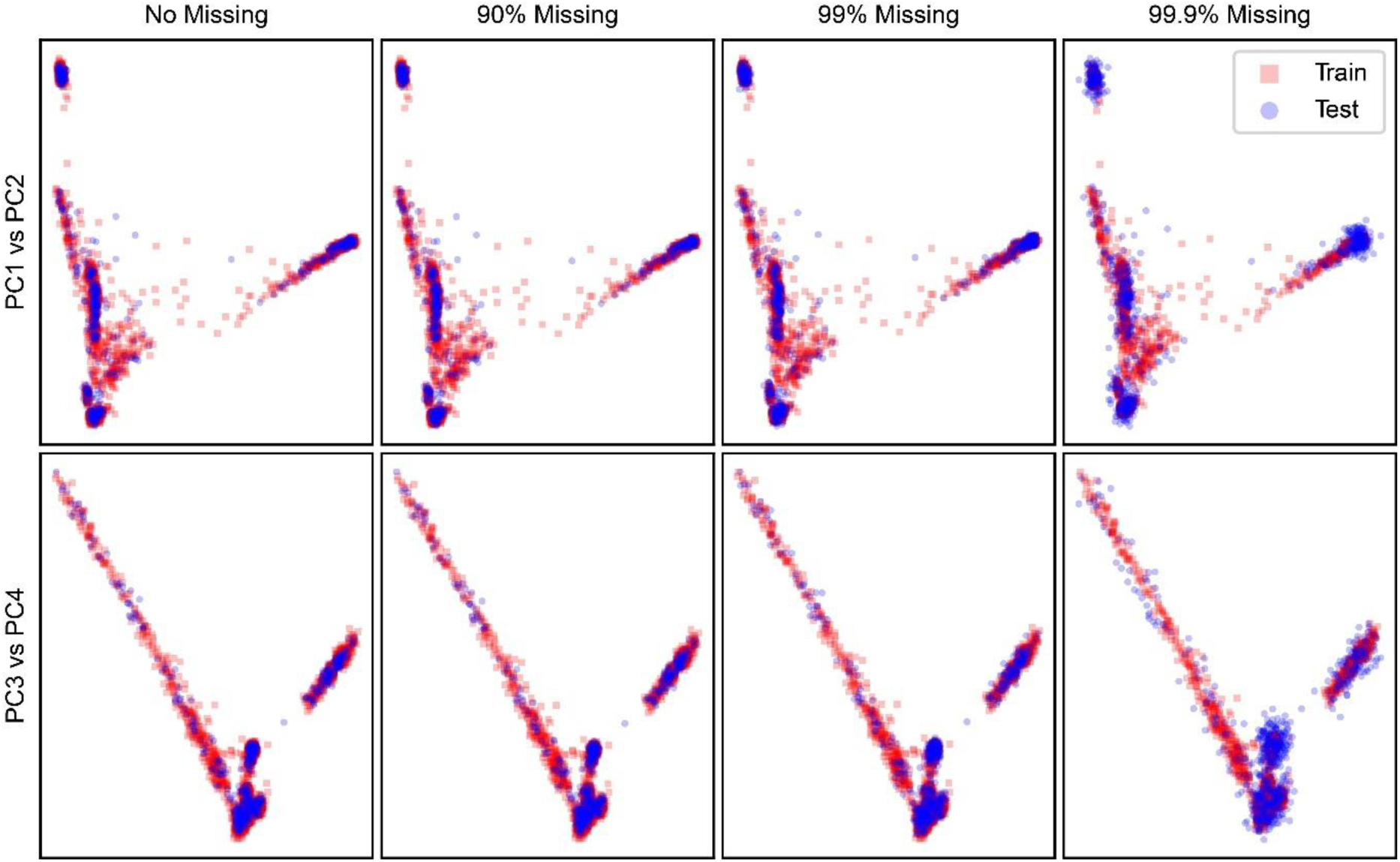
Scatter plots of PCA components of the 1KGP data. The top row is showing PC1 (horizontal) versus PC2 (vertical) and the bottom shows PC3 (horizontal) versus PC4 (vertical). Each column shows different test data with various degree of missing sites. The PCs with missing data were scaled by the inverse proportion of observed data. With higher degree of missing, the PCA components of the test data deviate more from that of the training data.

**Supplementary Figure 3.**
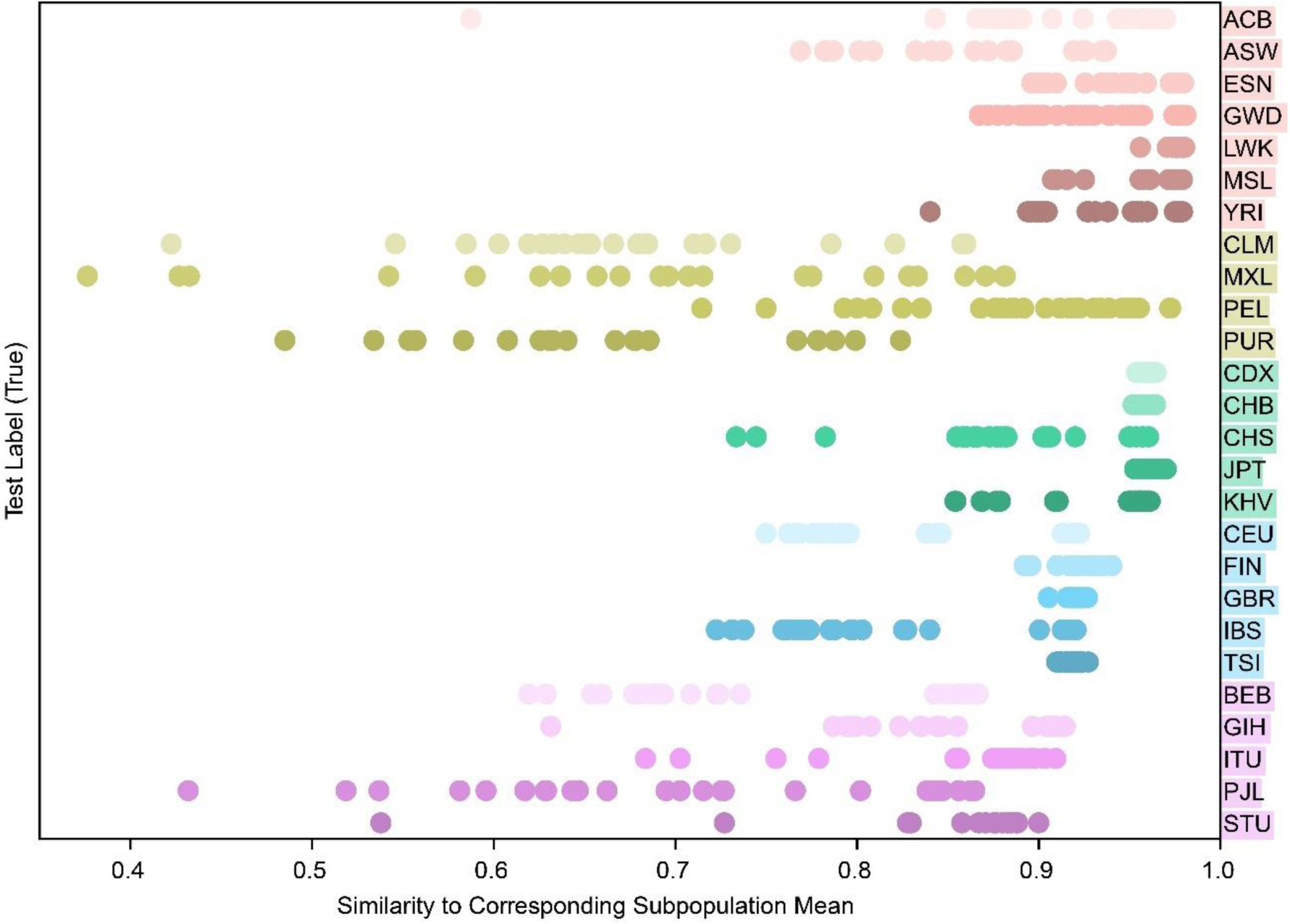
Heterogeneity within each population group. After splitting the 1KGP data into training-test (80-20%) and using the training data to construct the principal component space, we computed the cosine of the product of test data’s PC to its corresponding population mean PC vector. This computation captures how much each test individual deviates from the mean of its corresponding population group, and thereby the degree of heterogeneity within each population group. Some population groups, such as the Japanese, Japan (JPT) exhibit a low level of heterogeneity, whereas others, such as the Mexican Ancestry in LA, California (MXL) exhibit relatively higher levels of heterogeneity.

**Supplementary Figure 4.**
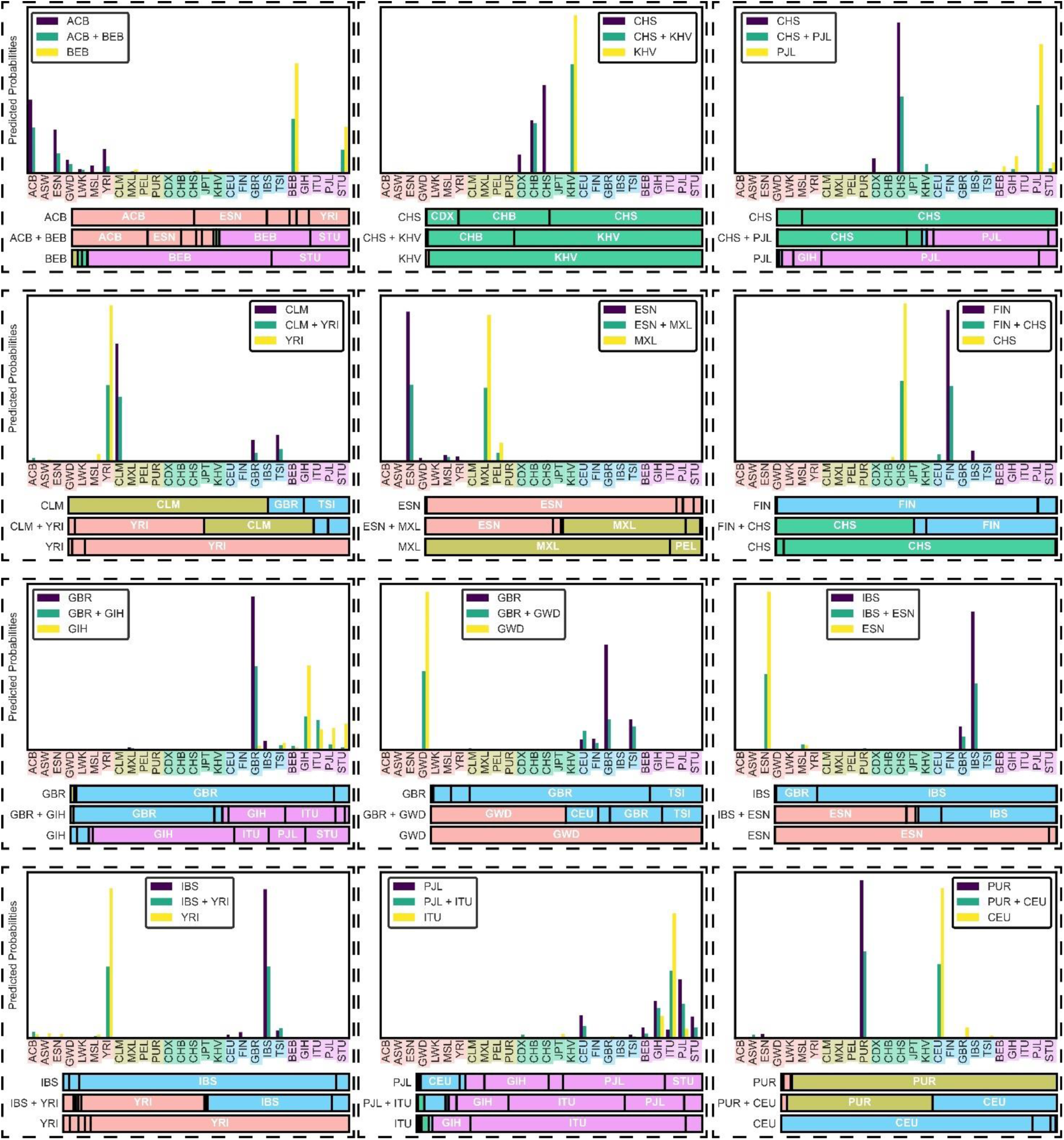
Above are 12 simulated trios of admixed individuals and their parents. For each plot, random individuals from two populations in the 1KGP datasets were selected (parents) and their genotypes were randomly shuffled to generate a new genotype vector that has 50% of each parent’s genotypes. This simulated admixed genotype vector was then used in the scAI-SNP model to get the inferred ancestry. As can be seen in both vertical and horizontal bar plots, the inferred ancestry of the admixed individual is almost a perfect mixture of the ancestry of the parents, showing that the model also works well with admixed individuals.

## Notes

https://gitlab.com/hormozlab/scAI-SNP

### Competing Interest Statement

The authors have declared no competing interest.

